# Isoleucine and asparagine influence the growth of staphylococcal strains in atopic dermatitis

**DOI:** 10.64898/2026.09.15.751654

**Authors:** Hogan Kok-Fung Wai, Xiaoye Wang, Clement Ching Yin Li, Gilbert T. Chua, Christina Sze Man Wong, Jaime S. Rosa Duque, Vivian Wai Yan Chan, Adrien Le Guennec, Jie Zhu, Shabnan Khan, Aina Navaz Punet, Mengzhen Ji, Jingmian Zhang, Elle Simone Hill, You Che, Helen Alexander, Eric Deharo, Hein Min Tun, David L. Moyes

## Abstract

Atopic dermatitis (AD) is a multifactorial, chronic skin disease associated with microbial factors. It is known that staphylococcal skin microbes play a role in the pathophysiology of AD, notably *S. aureus*. However, how staphylococcal strain variation affects disease aetiology and progression remains unclear. Using a culture-dependent approach, we analyzed the genomic and phenotypic profile of staphylococcal strains isolated from a Hong Kong children cohort (n = 70). Presence of *S. aureus* was positively associated with AD skin, whilst *S. epidermidis* and *S. hominis* were positively associated with healthy skin, but no overall strain type was associated with AD. In vitro experimentation revealed that there is demarcation in *S. aureus* strain growth, biofilm biomass formation, and metabolite production by disease severity. While isoleucine and asparagine promoted *S. aureus* growth, these amino acids suppressed *S. hominis* growth. These collective findings highlight the importance of strain-level and metabolic interactions of staphylococci in AD pathophysiology.

## Introduction

Atopic dermatitis (AD) is a chronic inflammatory skin disorder that is characterized by skin itch, crusted lesions, and lichenification ^1^. This disease is a global problem with a large epidemiological variation, affecting people of different ages, genders, and ethnicities. Approximately 2.6% of the global population is affected by AD, with the prevalence of AD having increased in the past few decades ^1–3^. Genetic, environmental, and microbial factors have all been shown to be risk factors for AD ^4^. As a result, developing an effective treatment for AD is challenging, as the pathophysiology of AD is complex and multifactorial.

Increasingly, skin microbes have been shown to play a role in AD. The role that the skin microbiome plays in the onset of AD starts from early life, as the skin microbiome is different between infants who developed AD in comparison to infants who did not ^5,6^. Later in life, this dysbiosis is still observed, with a lower alpha diversity on the skin of children and adults with AD as compared to healthy controls ^7,8^. *S. aureus* is a major player in the dysbiosis of the AD skin microbiome and has been associated with AD flares ^9^. This bacterium is more likely to colonize lesional skin of AD subjects as compared to non-lesional skin ^10^. Interestingly, the AD skin metabolome in lesions and samples with high *S. aureus* abundance have been shown to have increased branched chain amino acids production; this may provide a hypothesis for the metabolic effects of *S. aureus* in AD ^11^. Although *S. aureus* and its abundance is positively associated with AD, it is unclear why some patients have mild AD and others have severe AD, given its presence in most cases. The fact that no single strain of *S. aureus* is consistently associated with AD suggests that strain variation may play a role in disease heterogeneity ^12–18^.

Besides *S. aureus*, other *Staphylococcus* species, including *S. epidermidis* and *S. hominis,* play a role in AD. Both *S. epidermidis* and *S. hominis* have largely been shown to be protective in AD through multiple mechanisms. These include production of glutamyl endopeptidases and phenol soluble modulins by *S. epidermidis* to inhibit *S. aureus,* and inhibition of *S. aureus* psmα toxin production and agr quorum sensing by *S. hominis* ^19–23^. However, paradoxically *S. epidermidis* can also act as a pathogen in AD by damaging the skin barrier through secretion of an extracellular cysteine protease EcpA ^24^. There is also emerging evidence that *S. hominis* isolated from AD patients behaves differently from *S. hominis* isolated from control subjects by worsening cell viability and barrier function in vitro ^25^. The observation of the dual role that *S. epidermidis* and *S. hominis* plays in AD indicates the importance of strain heterogeneity and complexity for host-microbe interactions in disease.

The collective evidence suggests that a greater understanding of staphylococcal strain variation is necessary to discern its effect on disease severity and outcome. It remains unclear how different *Staphylococcus* strains contribute to AD. Notably, it is unknown whether there are differences in species composition and strains across different disease severities, and what the underlying mechanisms are in these differences. This study leverages the findings from a paediatric and adolescent cohort in Hong Kong (HK) with genomic and phenotypic experimental approaches to answer these questions.

## Methods

### Clinical study design

Between November 2021 and September 2022, a clinical case-control study was carried out with two study groups: AD subjects and non-atopic healthy controls (HC). In line with ethical requirements, the study was approved by the Institutional Review Board (IRB) of the HKU/ Hospital Authority HK West Cluster (IRB reference number: UW 21-559). All participants in this study were briefed about the study and provided written consent prior to enrolment.

### Participant recruitment

AD subjects were recruited from the Queen Mary Hospital in HK and an open recruitment call to the HKU population. Subjects recruited from the hospital had a diagnosis of AD based on the Hanifin and Rajka criteria ^26^. For subjects recruited from the open call at HKU, subjects were asked to confirm that they had been previously diagnosed by a clinician as having AD. All subjects diagnosed with AD had their disease severity determined using the SCORAD index ^27^. Non-atopic HC subjects were recruited from an open recruitment call to the HKU population. In both arms of the study, potential participants were excluded if they met one or more of the following criteria: (a) had a concurrent chronic inflammatory skin disease such as psoriasis and lichen planus; (b) had used systemic (oral or injection) corticosteroids in the past 4 weeks prior to sampling; (c) had used antibiotics in the past 4 weeks prior to sampling; (d) had used antifungal agents in the past 4 weeks prior to sampling; (e) had bathed or showered in the 6 hours prior to sampling; (f) was unable to cooperate with the study protocol for any reason at the time of sampling (Table S1).

### Sample collection

Sample collection was performed by trained research staff. Two areas of the subject’s body were targeted for sampling: one from a moist area of the body (i.e. antecubital fossa, popliteal fossa) and one from a corresponding dry area of the body (volar forearm, posterior thigh). If a subject did not have AD at the targeted site of sampling, then other areas of the body were selected instead. At each site of sampling, one swab was taken from a lesional area and one swab was taken from a non-lesional area. Therefore, a total of four swabs were taken from each subject. For HC subjects, the antecubital fossa, popliteal fossa, volar forearm, and posterior thigh were swabbed to enable matching with AD swabs. This was not adhered to only if it was known that a matching AD patient (by age and sex) was swabbed elsewhere on the body. In such a scenario, the HC subject would be swabbed according to the sites that the matching AD patient was sampled from. In total, four swabs were taken from each HC subject.

To collect a sample, an eSwab (Copan Diagnostics, Murrieta, CA) was premoistened with sterilized water and rubbed on the subject’s skin at the selected site for ten times back and forth (Han et al., 2018). Afterwards, the skin swabs were placed in a sterile tube containing stabilizer media (Liquid Amies Medium) (Copan Diagnostics, Murrieta, CA). Negative control swabs were also collected by waving premoistened swabs in the air for each clinical session. The samples were then temporarily stored in an ice bucket before being transported to the laboratory for processing and long-term storage in a -80°C freezer.

### Staphylococcus species isolation

Bacteria isolation was completed for each skin swab to isolate *Staphylococcus* species. The collected sample tube was pulse vortexed at high-speed for 10 s. For each sample, a total of 10 µL of the stabilizer media was inoculated in 9 mL of tryptone soy broth (TSB) (Oxoid, U.K.) and shaken overnight at 37°C and 220 rpm. A loopful of overnight culture for each sample was applied to a Brilliance Staph24 selective agar plate (Oxoid, U.K.) and incubated at 37°C. After an 18 h incubation period, up to five single colonies with unique bacteria morphology and colour were picked for each swab sample. Selected colonies were then individually grown overnight in TSB at 37°C and 220 rpm before storing the bacteria in 20% glycerol at -80°C.

### Bacterial species identification

Bacterial species identification was made using matrix-assisted laser desorption/ionization (MALDI-TOF) analysis. Cultures were analysed with the MALDI Biotyper® (Bruker, U.S.A.) using the flexControl 3.4 software and Biotyper 4.1.70 software with MBT version 7311 MPS library.

### Bacterial genomic DNA extraction

Bacterial genomic DNA was extracted from each sample using the QIAamp DNA Mini Kit (QIAGEN, Germany) following the manufacturer’s protocol. After the completion of DNA extraction, the amount of genomic DNA was quantified using the NanoDropND-1000 spectrophotometer (NanoDrop Technologies, U.S.A.).

### Whole genome sequencing

The bacterial genomic DNA samples were sent to Novogene (HK) Company Limited for whole genome sequencing. The NEBNext® Ultra™ IIDNA Library Prep Kit was used for library construction and microbial whole genome sequencing was performed on the Illumina NovaSeq PE150 platform with a 150 bp paired-end protocol.

### Whole genome assembly

Initially, a quality control check was completed on the short reads with FastQC v.0.12.1 and viewed with MultiQC v.1.30 ^28^. Next, the pair-end reads for all samples were assembled using Shovill v.1.1.0 (). After the genomes were assembled, the lineage specific workflow of CheckM v1.2.3 was used to calculate the completeness and contamination of assembled genomes ^29^. QUAST v5.3.0 was used to calculate the N50 and the number of contigs ^30^. The taxonomic classifications of assembled genomes were assigned using GTDB-Tk v2.1.055 based on the Genome Database Taxonomy GTDB (R07-RS207) ^31^. After checking and based on established quality control threshold used in the literature, isolates with draft genomes that had a completeness of > 95%, contamination < 5%, N50 > 50,000, and number of contigs < 100 were used for downstream processing ^16,32^.

### Staphylococcal protein A (spa) typing

The *spa* type of each *S. aureus* isolate was determined by amplifying the polymorphic X region of the *spa* gene with the forward primer 1095F (5’-AGAC GATCCTTCGGTGAGC-3’) and reverse primer 1517R (5’-GCTTTTGC AATGTCATTTACTG-3’) ^33^. PCR reaction volumes and conditions were conducted as previously benchmarked ^34^. The PCR products were purified using the QIAQuick PCR Purification Kit (QIAGEN, Germany) following the manufacturer’s instructions. After purification, the samples were sent to BGI Genomics HK for Sanger sequencing using the same primers for amplifying the *spa* gene.

Post sequencing, the *spa* types were determined using the *S. aureus spa* typing Task Template in the Bruker MBioSEQ™ Ridom Typer software version 10.5 (Bruker, U.S.A.). For *spa* types having fewer than five repeats, these were removed from clustering. The Based Upon Repeat Pattern (BURP) clustering algorithm was then used to group the assigned *spa* types into *spa*-clonal clusters (*spa*-CCs) if the cost between two different *spa* types was less than or equal to four ^35^. A *spa*-CC was created only if there were two or more related spa types in a cluster.

### Multilocus sequence typing

The *spa* type for each typable *S. aureus* isolates from Sanger sequencing reads was mapped to a MLST ST using the Ridom spa server (https://www.spaserver.ridom.de/; date accessed – June 2025). A *spa* type that was not mapped to a MLST ST was marked as having no predicted MLST ST. Subsequently, each mapped MLST ST was then associated with a MLST CC based on a literature search. Mapping was only possible if there were past studies that have done both *spa* typing and MLST PCRs. To ensure unambiguous mapping, the mapped MLST STs that have been associated with more than one ST in the literature were not associated with a MLST CC.

Those *S. aureus* isolates with a WGS assembly were typed with MLST in silico using the *S. aureus* MLST Task Template in the Bruker MBioSEQ™ Ridom Typer software version 10.5 (Bruker, U.S.A.). *S. hominis* isolates were typed with MLST in silico. This was done using the *S. hominis* MLST Task Template in the Bruker MBioSEQ™ Ridom Typer software version 10.5 (Bruker, U.S.A.). Novel MLST STs were submitted to the PubMLST database (https://pubmlst.org/) for curation. Following the assignment of a MLST ST for an isolate, the eBURST algorithm within the Bruker MBioSEQ™ Ridom Typer software version 10.5 (Bruker, U.S.A.) was used to cluster the MLST typed strains. For each species, a MLST-CC was formed if the STs for matched the founder ST at four or more loci. The founder ST was defined to be the ST that had the highest number of single-locus variants.

### Growth curves assay

The growth curves assay was used to generate growth curves for each *Staphylococcus* strain. 1 µL of 1:10^8^ dilution of overnight culture in PBS was added to 99 µL of TSB in each well of a sterile 96-well Microplate. Isoleucine and asparagine were sourced from Thermo Scientific U.K. and Sigma Aldrich U.K., respectively. The sealed plate was placed into the Multiskan™ FC Microplate Photometer (Thermo Scientific, U.K.) for simultaneous incubation at 37°C and shaking at slow speed (5 Hz, amplitude 15 mm). Readings were taken every 15 min at 620 nm for 24 h. To obtain the growth rate (r) and carrying capacity (k) from each growth curve, the Growthcurver R package was used ^36^. The clustering of the strains by r and k was done using the k-means clustering method, with the optimal number of clusters selected using the elbow method ^37^.

### Biofilm culturing

Overnight cultures were diluted to an OD_600_ of 0.5 in 1X PBS. Subsequently, 100 µL of each sample was pipetted into an individual well of a 24-well plate (Thermo Scientific, U.K.) on ice. Wells were topped up with 400 µL of TSB and incubated for 24 h at 37°C. Media was replenished every 24 h for 2 days.

### Biofilm crystal violet assay

Biofilm biomass was quantified using the crystal violet assay. The biofilm was washed once with PBS before being fixed with 300 µL of methanol and stained with 300 µL of 1% crystal violet solution (Sigma Aldrich, U.K.) for 15 min and then rinsed with deionised water. The stained biofilm was then solubilized using 300 µL of 30% acetic acid (Sigma Aldrich, U.K.). A 100 µL aliquot was transferred to a 96-well Microplate and the absorbance was measured at 620 nm.

### Biofilm XTT assay

The sodium 2,3-Bis-(2-Methoxy-4-Nitro-5-Sulfophenyl)-2H-Tetrazolium-5-Carboxanilide (XTT) assay was used to measure biofilm metabolic activity using the Cell Proliferation Kit II (Roche, Germany) according to manufacturer’s protocol. The biofilm was washed once with PBS before 400 µL of TSB and 100 µL of XTT reagent were added. A negative control well was also included to control for background absorbance. The biofilms were incubated for 4 h, after which a 100 µL aliquot was transferred to a Nunc™ MicroWell™ 96-well Microplate (Thermo Scientific, U.K.) and the plate was read once at an absorbance of 450 nm and 689 nm with the Multiskan™ FC Microplate Photometer (Thermo Scientific, U.K.). The final absorbance value for each sample was calculated by taking the difference between the blanked 450 nm reading and blanked 689 nm reading for each sample.

### NMR metabolomics

Nuclear magnetic resonance (NMR) metabolomics was performed at the King’s College London NMR Facility to identify metabolites produced. Each strain was initially grown for 18h at 37°C and 220 rpm. After culturing, the samples were centrifuged for 10 min at 4,000 x g. Supernatants were then filtered through a sterile 0.2 µM syringe filter (Fisher Scientific, U.K.). For each sample, a ^1^H spectra was generated using a 600 MHz NMR spectrometer and the peaks in the spectra were picked. Intensity values for a metabolite that were 1.5 * interquartile range greater than the third quartile or 1.5 * interquartile range less than the first quartile were classified as outliers and removed from the dataset.

### Statistical analyses

For comparing distributions or detecting potential associations, Spearman’s correlation with t-test, Kruskal-Wallis test (with Dunn’s post hoc test for multiple comparisons), and Fisher’s exact test (with false discovery correction for multiple comparisons) were used. The Phi coefficient (Φ) was used to assess the strength and direction of association between two binary variables, with the Fisher’s exact test used to determine if the calculated coefficient was statistically significant. All statistical analyses were performed using the RStudio software (version 2024.12.1+563).

## Results

### S. aureus is positively associated with AD, while S. hominis is negatively associated with AD

To determine staphylococcal species diversity, a clinical cohort of 70 participants (46 AD and 24 HC) were recruited from HK (Figure 1A, Table 1). A total of 835 staphylococcal isolates were cultured from the skin of the participants. The cultured species included *S. aureus* (340/835; 41%), *S. capitis* (111/835; 13.3%), *S. epidermidis* (144/835; 17.2%), and *S. hominis* (97/835; 11.6%) (Table S2). Of these, *S. hominis* and *S. epidermidis* were predominantly isolated from healthy individuals as compared to AD patients (q < 0.001 and q < 0.01, respectively) (Table S3, Table S4). In contrast, *S. aureus* was the major *Staphylococcus* species colonizing the skin of diseased individuals (q < 0.001) (Table S3, Table S4).

**Figure 1.**
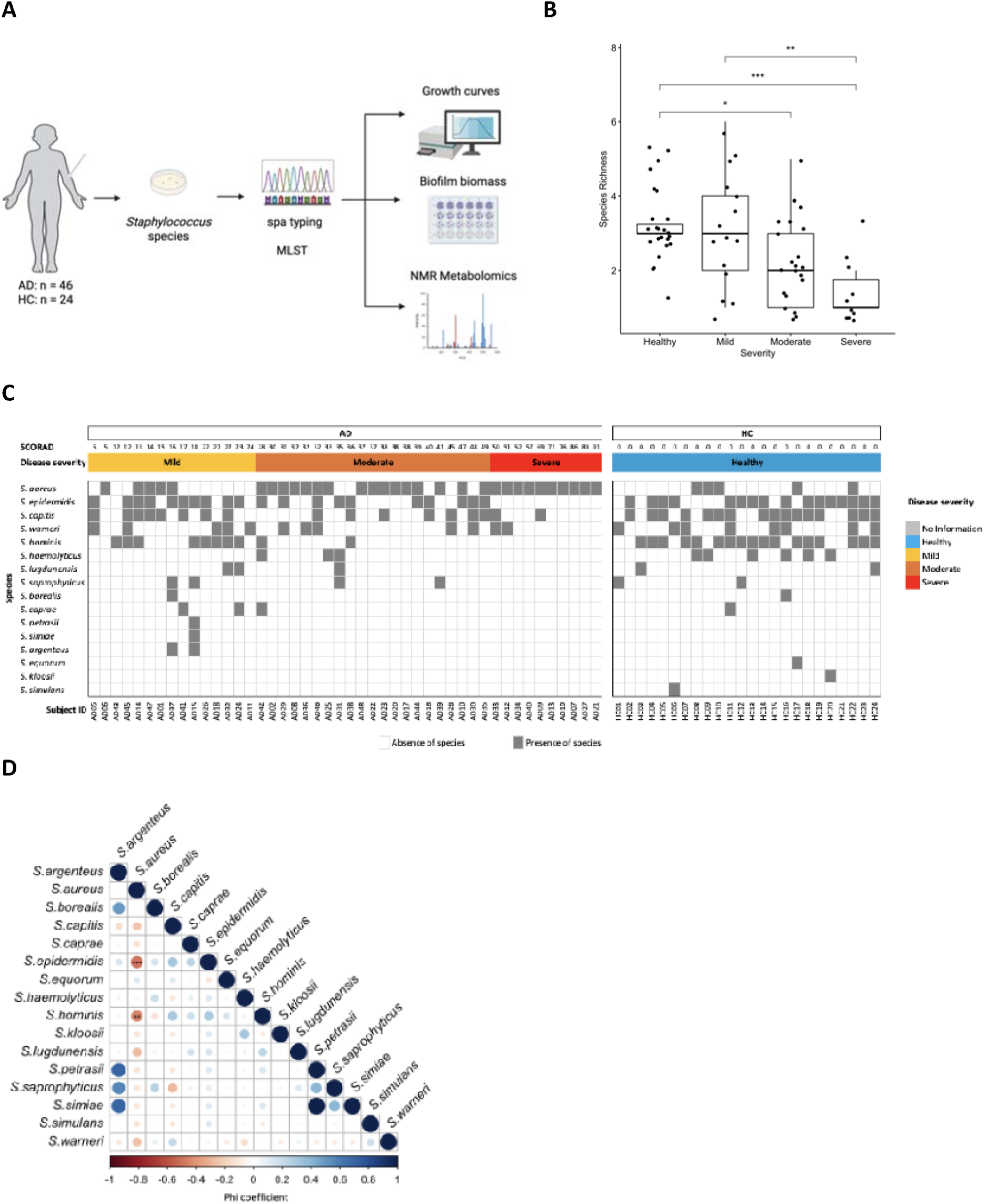
Concurrent S. aureus colonization and Staphylococcal diversity reduction is a hallmark of AD. (A) Diagram of study design. (B) Boxplot of the *Staphylococcus* species richness by severity. Each datapoint represents the number of *Staphylococcus* species that could be isolated from one subject. Significance levels are denoted as follows: * p < 0.05; ** p < 0.01; *** p < 0.001 (by Dunn’s test). (C) Stacked tile plot of the absence and presence of *Staphylococcus* species in 46 AD and 24 HC subjects. (D) Pairwise association matrix showing the cooccurrence of *Staphylococcus* species for 46 AD subjects and 24 HC subjects. Each circle represents the Phi coefficient for each pair of species. The Phi coefficient ranges from -1 to +1, where -1 (in dark red) indicates a perfect negative association (two species never occur together; co-absence), 0 (in white) indicates no association (mutual exclusion), and +1 (in dark blue) indicates a perfect positive association (two species always occur together; co-presence). The size of each circle shows the absolute value of corresponding Phi coefficients. Significance levels are denoted as follows: ** q < 0.01; *** q < 0.001 (by Fisher’s test followed by FDR-correction).

**Table 1.** Participant demographics.

| Characteristic | AD<br>N = 46<sup>1</sup> | HC<br>N = 24<sup>1</sup> | p-value<sup>2</sup> |
| --- | --- | --- | --- |
| Age | 12 (4, 15) | 5 (3, 9) | 0.015 |
| Gender |  |  | 0.9 |
| Female | 20 (43%) | 11 (46%) |  |
| Male | 26 (57%) | 13 (54%) |  |
| Severity |  |  | <0.001 |
| Healthy | 0 (0%) | 24 (100%) |  |
| Mild | 15 (33%) | 0 (0%) |  |
| Moderate | 21 (46%) | 0 (0%) |  |
| Severe | 10 (22%) | 0 (0%) |  |

<sup>1</sup> Median (Q1, Q3); n (%)
<sup>2</sup> Wilcoxon rank sum test; Pearson's Chi-squared test; Fisher's exact test

### Staphylococcal species richness is associated with disease severity

The distribution of *Staphylococcus* species isolated in the study participants varied depending on disease status (Figure 1C). The mean number of unique *Staphylococcus* species that could be isolated from subjects decreased with increasing disease severity: 3 species from healthy subjects, 3 species from mild AD patients, 2 species from moderate AD patients and 1 species from severe patients (Figure 1B). Notably, there was a significant difference in the mean number of unique *Staphylococcus* species isolated from healthy participants as compared to severe AD patients (p < 0.001).

To build upon these findings, a pairwise association matrix was constructed to visualize the cooccurrence of different *Staphylococcus* species with one another (Figure 1D). Excluding pairing between the same *Staphylococcus* species, there were a total of 120 possible combinations of *Staphylococcus* species cooccurring with each other. Out of these, 14 combinations had a moderate-to-strong association of cooccurrence (Φ ≥ 0.3). Of these combinations only two pairings were statistically significant and inversely associated: *S. aureus*-*S. hominis* (Φ = -0.5; q < 0.01) and *S. aureus*-*S. epidermidis* (Φ = -0.45; q < 0.001).

### No single S. aureus strain was associated with AD

To determine whether there were associations between particular strains of *S. aureus* and disease severity, isolated strains were typed with a cost effective and rapid method: *spa* typing. The *spa* typing analysis revealed 29 distinct *spa* types among the 340 *S. aureus* isolates that successfully underwent *spa* typing (Table S5). The top five most common *spa* types were: t084 (62/340; 18%), t127 (44/340; 13%), t091 (37/340; 11%), t085 (22/340; 6%), and t189 (20/340; 6%) (Figure 2A). Outside of these top five *spa* types, none of the remaining *spa* types comprised more than 5% of the typed *S. aureus* isolates (Figure 2A, Table S5).

**Figure 2.**
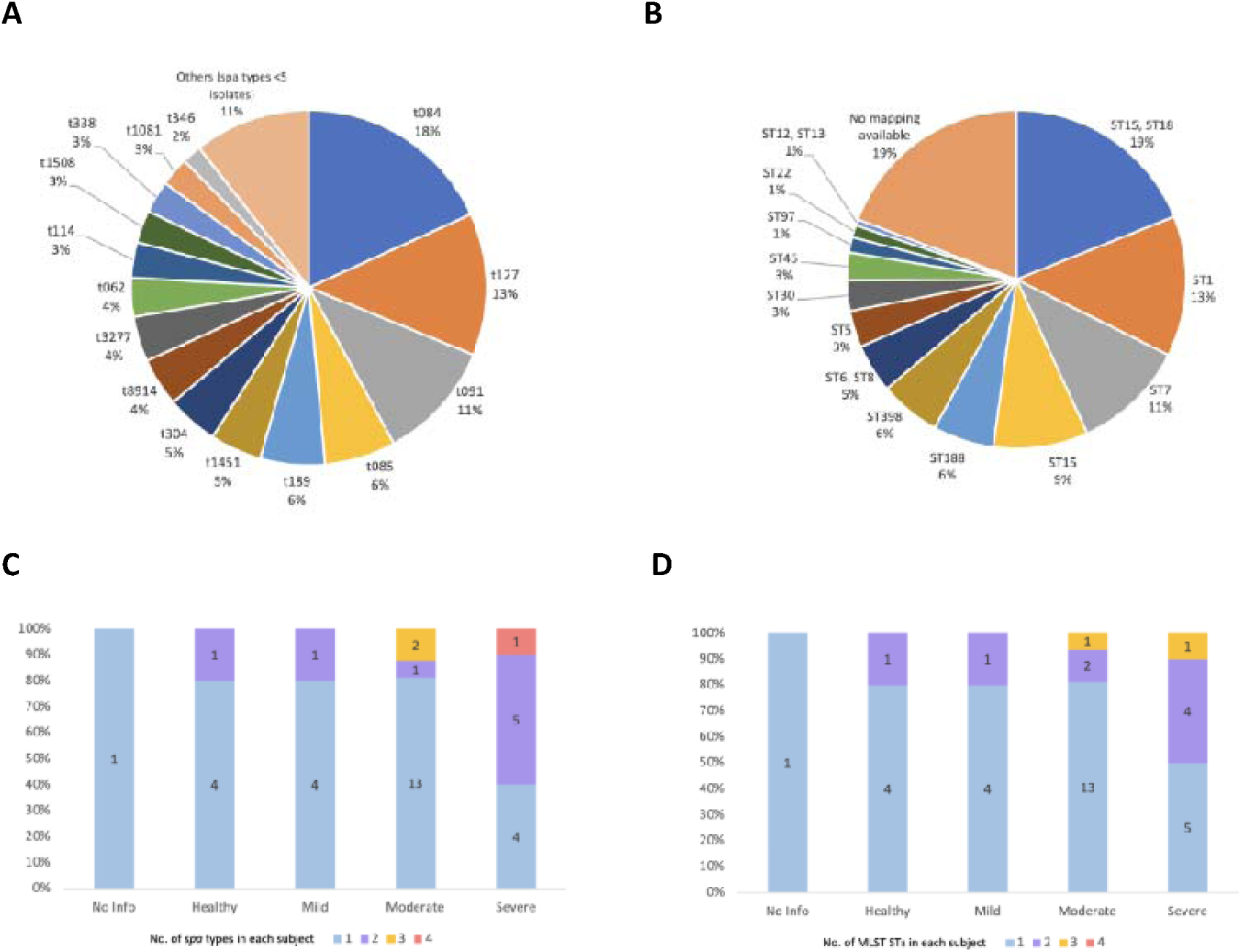
Severe AD is associated with a greater number of S. aureus genotypes found in a patient. (A) Distribution of *S. aureus* isolates grouped by spa type. (B) Distribution of *S. aureus* isolates grouped by mapped MLST ST. (C) Breakdown of the number of *spa* types found in each subject by severity. The colour of the bars indicates the number of *spa* types found in each subject. Data labels within each stacked bar show the number of subjects. (D) Breakdown of the number of MLST STs found in each subject by severity. The colour of the bars indicates the number of MLST STs found in each subject. Data labels within each stacked bar show the number of subjects.

Excluding *spa* types with less than 5 repeats (n = 2), the remaining identified *spa* types (n = 338) were then clustered into *spa*-CCs by BURP analysis. These five *spa*-CCs were: *spa*-CC 084, *spa*-CC 127, *spa*-CC 189, *spa*-CC 1451, and *spa*-CC 8914 (Figure S1, Table S5). Of the *spa* types isolated, 60% belonged to either spa-CC 084 (138/340; 41%) and spa-CC 127 (65/340; 19%) (Table S5). *spa*-CC 084 contained the highest number of *spa* types, with 9 *spa* types belonging to this CC. For the other four *spa*-CCs, there were no more than 3 *spa* types within each *spa*-CC. There were 9 *spa* types (60/340; 18%) that did not cluster into any *spa*-CCs and were classified as singletons (Table S5).

As identified *spa* types have been used to infer MLST STs reliably in previous studies, the obtained *spa* types were next mapped to a MLST ST ^33,38^. In this study, the typed *S. aureus* isolates generally mapped to a single MLST-ST (274/340; 81%) (Figure 2B, Table S5). Four MLST STs made up about half (51%) of the STs mapped: ST15/ST18 (62/340; 18%), ST1 (44/340; 13%), ST7 (37/340; 11%), and ST15 (31/340; 9%) (Figure 2B, Table S5). The other mapped MLST STs individually did not make up more than 6% of all MLST STs. For *spa* types that were not mapped to a MLST ST (66/340; 19%), this was due to the unavailability of such a mapping.

After mapping the *spa* types to MLST STs, we next inferred MLST-CCs from these MLST STs. There were 10 MLST-CCs inferred from the mapped MLST STs (Table S5). Over half of the strains (58%) were associated with one of three MLST-CCs: CC15 (96/340; 28%), CC1 (66/340; 19%), and CC7 (37/340; 11%) (Table S5). The 2 remaining MLST-CCs that made up the top 5 predicted MLST-CCs were: CC398 (20/340; 6%), and CC5 (12/340; 4%) (Table S5). Outside of these top five predicted MLST-CCs, none of the remaining associated MLST-CCs individually comprised more than 3% of the *S. aureus* isolates. With one of the goals of this study being to understand if there were any genotypic signatures associated with clinical phenotypes, we then tested the association between the compiled *spa* types and MLST STs with severity status, but found no significant associations (Table S6, Table S7). Similarly, there was no significant relationship between *spa*-CCs and AD severity and between MLST-CCs and AD severity (Table S8, Table S9).

### Number of S. aureus strains is associated with disease severity

While there were no specific *S. aureus* strains associated with disease, analyzing the number of *S. aureus* strains isolated from a participant revealed that 70% of participants were colonized with only 1 *spa* type (26/37) (Figure 2C). However, in the severe group a significant proportion of patients had ≥2 *spa* types (6/10; 60%) (Figure 2C). For patients that had 3 or more unique *spa* types on their skin, the *spa* types found in each patient belonged to the same *spa*-CC. These were: AD21 (colonized with *spa* types from *spa*-CC 127: t127, t386, t3887, and t6297), AD22 (colonized with *spa* types from *spa*-CC 084: t091, t803, t2119), and AD23 (colonized with *spa* types from *spa*-CC 084: t084, t346, t7214) (Figure S1). When the analysis of the number of *spa* types found in each individual and its association with disease severity was repeated for MLST ST, we also found that the number of strains was associated with disease severity.

### S. aureus strains from moderate and severe AD grew faster but to a lower density

For subsequent phenotypic analysis, a subset of isolates from the collection of *Staphylococcus* strains was picked. We selected isolates from *S. aureus* and *S. hominis* as these species were shown to be positively and negatively associated with AD, respectively (Table S3, Table S4). We included *S. hominis* as while *S. epidermidis* was also negatively associated with AD, it has previously been identified as a virulence factor in AD ^24^. The *S. aureus* and *S. hominis* isolates were further narrowed down based on strain type and severity status (Table S10, Table S11). After screening, 28 *S. aureus* isolates (6 isolates from healthy, 3 isolates from mild, 11 isolates from moderate, and 8 isolates from severe) and 12 *S. hominis* isolates (5 isolates from healthy, 5 isolates from mild, and 2 isolates from severe) were selected for onward phenotypic analysis.

To determine the growth rate and carrying capacity for the different strains, 24 h growth curves were generated for each isolate. To visualize the relationship between the growth curves of these strains, the mean growth rate (*r*) and carrying capacity (*k*) of the individual growth curves for each strain were determined (Figure 3A). For *S. aureus* strains, there was a significant negative Spearman correlation between the *r* and *k* (p < 0.01) (Figure 3A), indicating faster growing strains had a lower carrying capacity, while slower growing strains had a higher carrying capacity. These slower growing strains came from more severe AD disease (i.e. moderate or severe) and clustered together (Figure 3A). For *S. hominis*, there was no significant Spearman correlation between *r* and *k* of the isolated strains (Figure 3B). Moreover, the *S. hominis* strains showed no observable pattern of clustering based on *r* and *k*.

**Figure 3.**
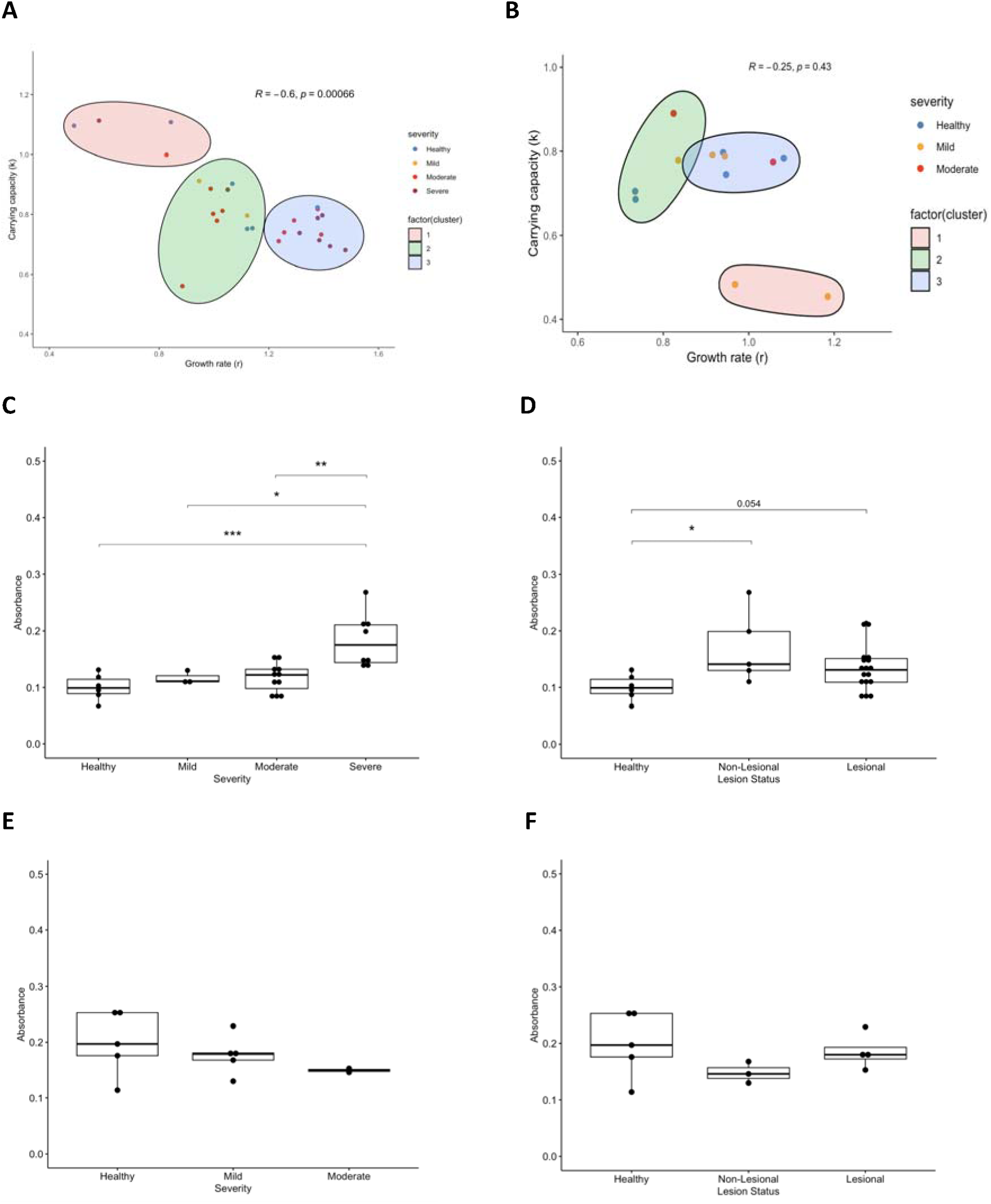
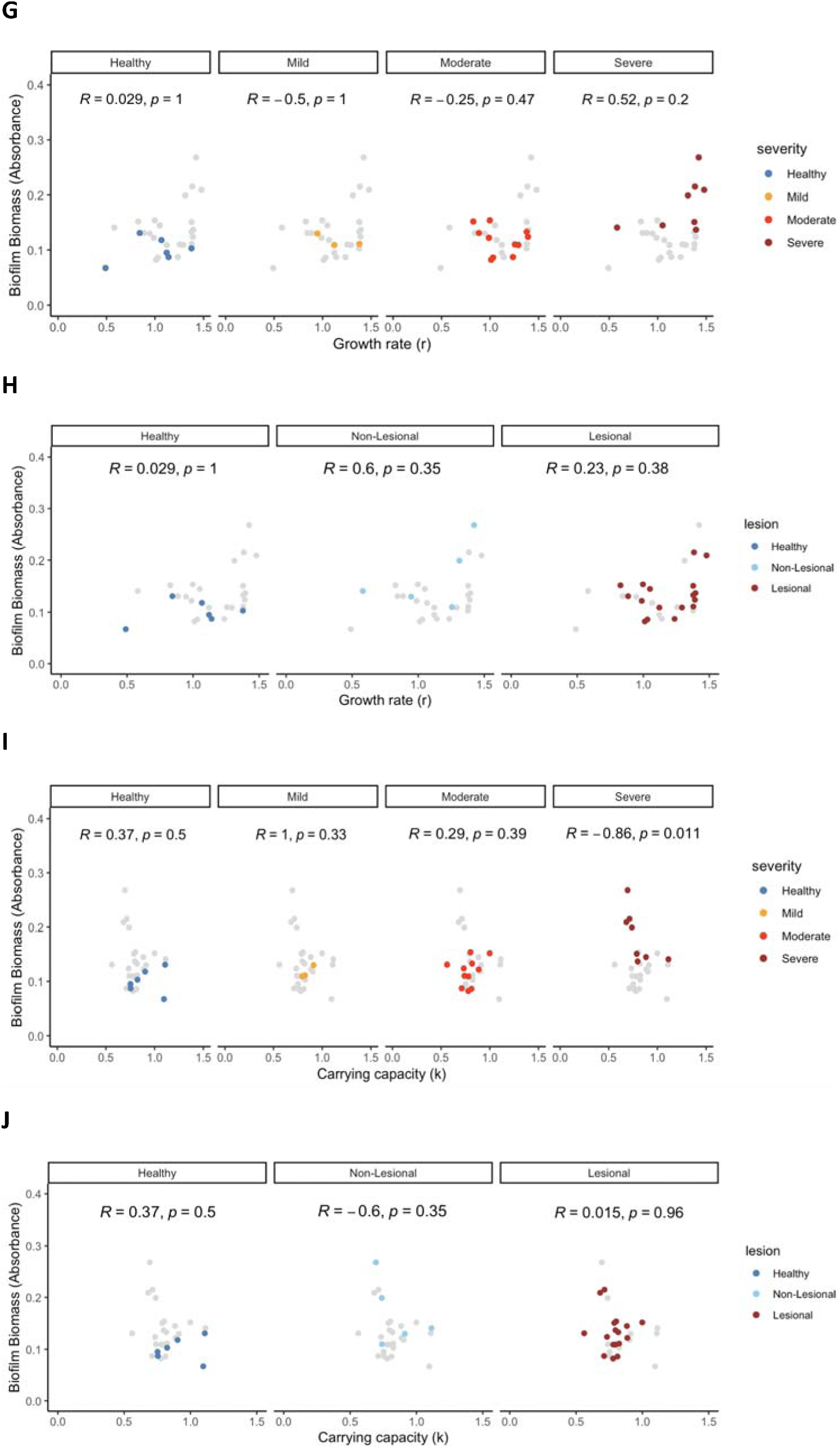
Staphylococcal growth and biofilm formation are correlated with severity and lesion status. (A) Association of growth rate (*r*) and carrying capacity (*k*) for *S. aureus* strains. *S. aureus* clinical strains were individually grown in TSB for 24 h and each data point shown in the graph represents *r* and *k* for each strain. Data points shown are the mean of three independent experiments. Clusters by severity were defined by k-means unsupervised clustering. Statistical significance of the Spearman correlation was determined using the t-test. (B) Association of growth rate (*r*) and carrying capacity (*k*) for *S. hominis* strains. *S. hominis* clinical strains were individually grown in TSB for 24 h and each data point shown in the graph represents *r* and *k* for each strain. Data points shown are the mean of three independent experiments. Clusters by severity were defined by k-means unsupervised clustering. Statistical significance of the Spearman correlation was determined using the t-test. (C) Biofilm biomass of isolated *S. aureus* strains by disease severity. Data points shown are the mean of three independent experiments. Significance levels are denoted as follows: * p < 0.05; ** p < 0.01; *** p < 0.001 (by Dunn’s test). (D) Biofilm biomass formation of isolated *S. aureus* strains by lesional status. Data points shown are the mean of three independent experiments. Significance levels are denoted by: * p < 0.05 (by Dunn’s test). (E) Biofilm biomass formation of isolated *S. hominis* strains by disease severity. Data points shown are the mean of three independent experiments. Statistical significance was determined by Dunn’s test. (F) Biofilm biomass of isolated *S. hominis* strains by lesional status. Data points shown are the mean of three independent experiments. Statistical significance was determined by Dunn’s test. (G) The association of *S. aureus* strain growth with biofilm biomass formation by disease severity and growth rate (*r*). Each data point represents a *S. aureus* strain, with colored dots used to showing the strains by disease severity. Data points shown are the mean of three independent experiments. Statistical significance of the Spearman correlation was determined using t-test. (H) The association of *S. aureus* strain growth with biofilm biomass formation by lesional status and growth rate (*r*). Each data point represents a *S. aureus* strain, with colored dots used to showing the strains by lesional status. Data points shown are the mean of three independent experiments. Statistical significance of the Spearman correlation was determined using t-test. (I) The association of *S. aureus* strain growth with biofilm biomass formation by disease severity and carrying capacity (*k*). Each data point represents a *S. aureus* strain, with colored dots used to showing the strains by disease severity. Data points shown are the mean of three independent experiments. Statistical significance of the Spearman correlation was determined using t-test. (J) The association of *S. aureus* strain growth with biofilm biomass formation by lesional status and carrying capacity (*k*). Each data point represents a *S. aureus* strain, with colored dots used to showing the strains by lesional status. Data points shown are the mean of three independent experiments. Statistical significance of the Spearman correlation was determined using t-test.

### Biofilm formation in S. aureus strains is positively associated with severity and lesion status

The ability of microbes to form biofilms is also a key aspect of virulence ^39^. To determine if there were any disease associated strain variation in biofilm formation, the *S. aureus* and *S. hominis* strains were cultured as biofilms and their biomass measured. Across disease severity and lesion status, AD *S. aureus* strains were better able to form biofilms (Figure 3C). The *S. aureus* strains from the severe AD group generated more biofilm biomass as compared to the moderate AD group (p < 0.01), mild AD group (p < 0.05), and healthy group (p<0.001) (Figure 3C). Similarly, *S. aureus* strains isolated from both lesions (p = 0.054) and non-lesions (p < 0.05) of AD patients generated more biofilm biomass than strains from healthy controls (Figure 3D).

For *S. hominis* strains, there was no statistically significant difference in biofilm formation between strains clustered by disease severity (Figure 3E) or lesion status (Figure 3F). However, there is a trend showing *S. hominis* strains from healthy subjects were better able to form biofilm biomass as compared to strains from diseased subjects (Figure 3E). Similarly, *S. hominis* strains coming from healthy skin formed more biofilm biomass than strains from lesional or non-lesional on AD skin (Figure 3F).

### Biofilm formation in S. aureus strains is negatively associated with carrying capacity

Given that there were statistically significant differences in the growth (rate/carrying capacity) and biofilm biomass formation of *S. aureus* strains, we next determined if these two phenotypic parameters were correlated. There is a clustering of strains by disease severity and lesional status between *S. aureus* for growth rate and biofilm biomass formation (Figure G, Figure H). A similar clustering is seen when looking at carrying capacity and biofilm biomass formation (Figure 3I, Figure 3J). Notably, the carrying capacity and biofilm biomass formation of strains from severe AD patients were negatively correlated (*R* = -0.86; p = 0.011) (Figure 3I). However, this was not observed for clustering of the *S. aureus* strains by lesional status (Figure 3J), where no correlation is seen (*R* = 0.015; p = 0.96).

### Isoleucine and asparagine are produced in higher levels by disease derived S. aureus strains

As the *Staphylococcus* strains showed differences in growth and biofilm biomass formation, we postulated that it is likely that they would produce different levels of key metabolites. To measure the metabolome produced by each strain, exhausted media from overnight cultures was analysed using NMR. Overall, 28 metabolites were identified at intensity levels above background (Table S12). Of note, close to half of the metabolite types identified were amino acids (13/28; 46%). 18% of identified metabolites (5/28) were classified as carbohydrates/sugars (mannose, glucose, raffinose, and trehalose). A further 5 metabolites were classified as organic acids/ TCA cycle intermediates (pyruvate, fumaric acid, citrate, acetate, and lactate). Based on the list of metabolites identified, a comparison of the levels of the metabolites present between AD and HC was conducted to understand which metabolites were higher or lower by disease group.

While there were no statistically significant differences in the metabolites produced by *S. hominis* strains from AD and HC participants, there was a higher level of isoleucine (q < 0.05) and asparagine (q < 0.05) produced by *S. aureus* strains from AD as compared to HC (Table 2, Table S13). As compared to strains isolated from healthy participants, *S. aureus* strains from mild (p < 0.05), moderate (p < 0.01) and severe AD (p < 0.01) produced a higher level of isoleucine (Figure 4A). There was also a higher isoleucine level for *S. aureus* strains that were isolated from AD skin (lesional and non-lesional) (p < 0.01) as compared to HC skin (Figure 4B). However, there was no significant difference when comparing the levels of isoleucine between diseased *S. aureus* strains by AD severity and by lesion status (Figure 4A, Figure 4B). Based on this observation, it can be concluded that rather than disease severity or lesion status, it is the disease state (i.e. AD or HC) from which a *S. aureus* strain was isolated from that delineates the difference in isoleucine production.

**Figure 4.**
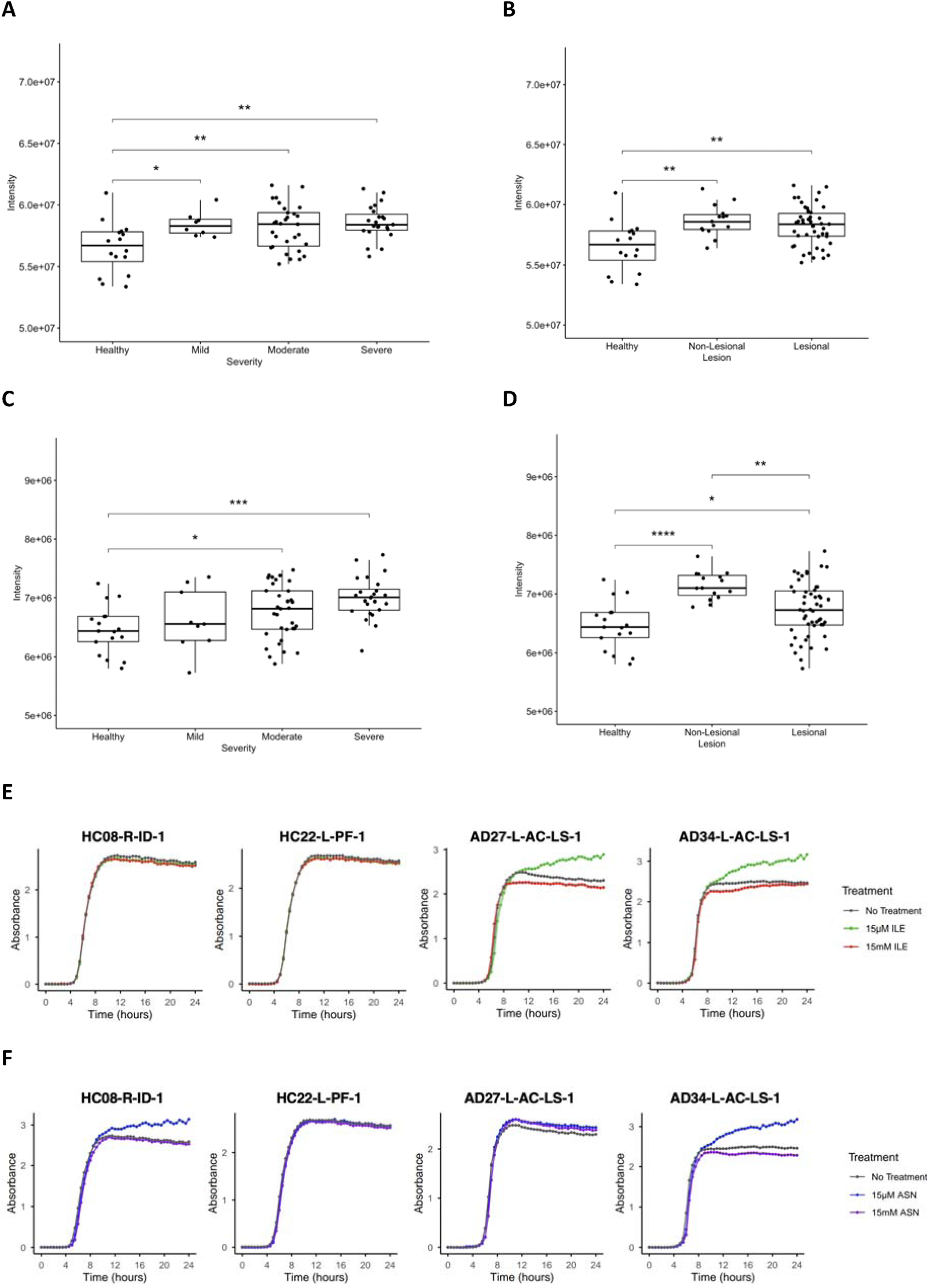
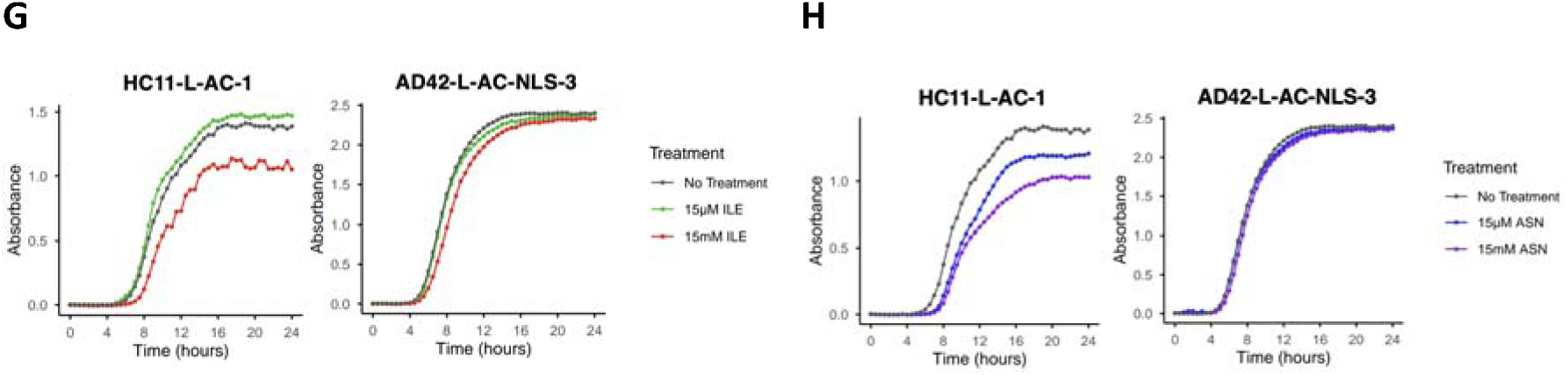
Isoleucine (ILE) and asparagine (ASN) can increase S. aureus growth, while reducing S. hominis growth. (A) Intensity levels of isoleucine produced by S. aureus strains shown by disease severity. Each data point represents the biological repeat for a S. aureus strain. Significance levels are denoted as follow: * p < 0.05; ** p < 0.01 (by Dunn’s test). (B) Intensity levels of isoleucine produced by S. aureus strains shown by lesion status. Each data point represents the biological repeat for a S. aureus strain. Significance levels are denoted as follow: ** p < 0.01 (by Dunn’s test). (C) Intensity levels of asparagine produced by S. aureus strains shown by disease severity. Each data point represents the biological repeat for a S. aureus strain. Significance levels are denoted as follow: * p < 0.05; *** p < 0.001 (by Dunn’s test). (D) Intensity levels of asparagine produced by S. aureus strains shown by lesion status. Each data point represents the biological repeat for a S. aureus strain. Significance levels are denoted as follow: * p < 0.05; *** p < 0.001; **** p < 0.0001 (by Dunn’s test). (E) 24 h growth curves of selected S. aureus strains with no isoleucine treatment, 15µM of isoleucine, and 15 mM of isoleucine. (F) 24 h growth curves of selected S. aureus strains with no asparagine treatment, 15µM of asparagine, and 15 mM of asparagine. (G) 24 h growth curves of selected S. hominis strains with no isoleucine treatment, 15µM of isoleucine, and 15 mM of isoleucine. (H) 24 h growth curves of selected S. hominis strains with no asparagine treatment, 15µM of asparagine, and 15 mM of asparagine.

**Table 2.** Distribution of metabolites produced by S. aureus strains from AD and HC.

| **Metabolite** | **p-value<sup>1</sup>** | **q-value<sup>2</sup>** |
| --- | --- | --- |
| Isoleucine | 0.001 | 0.031 |
| Asparagine | 0.003 | 0.042 |
| Ethanol | 0.014 | 0.137 |
| Lactate | 0.026 | 0.184 |
| Acetate | 0.032 | 0.184 |
| Valine | 0.048 | 0.231 |
| Pyruvate | 0.066 | 0.258 |
| Ornithine | 0.071 | 0.258 |
| Propylene glycol 2-3 Butandiol | 0.105 | 0.339 |
| Formate | 0.143 | 0.413 |
| Mannose | 0.185 | 0.489 |
| Leucine | 0.232 | 0.523 |
| Phenylalanine | 0.234 | 0.523 |
| Lysine | 0.275 | 0.54 |
| Orotate | 0.285 | 0.54 |
| Tryptophan | 0.298 | 0.54 |
| Tyrosine | 0.331 | 0.565 |
| Alanine | 0.386 | 0.608 |
| Glucose | 0.398 | 0.608 |
| Threonine | 0.463 | 0.656 |
| Uridine | 0.475 | 0.656 |
| Pyroglutamate | 0.503 | 0.662 |
| Methionine | 0.667 | 0.83 |
| Uracil | 0.744 | 0.83 |
| Fumaric acid | 0.745 | 0.83 |
| Citrate | 0.773 | 0.83 |
| Raffinose | 0.806 | 0.835 |
| Trehalose | 0.908 | 0.908 |

<sup>1</sup> Mann-Whitney U test
<sup>2</sup> q-value is the p-value with FDR correction

Similar to isoleucine, asparagine was produced at higher levels in *S. aureus* strains from AD as compared to HC participants (Figure 4C, Figure 4D). There was an increase in the level of asparagine produced by *S. aureus* strains that came from patients with more severe AD (p < 0.001) (Figure 4C). However, as with isoleucine, there was no significant difference when comparing the asparagine levels between *S. aureus* strains from AD patients by severity (Figure 4C). Asparagine levels produced by *S. aureus* strains that came from lesional and non-lesional areas of AD skin were statistically significant when compared with each other (p < 0.01) and with healthy skin (p < 0.05 and p < 0.0001, respectively) (Figure 4D).

### Isoleucine increases growth of S. aureus isolates from diseased but not healthy participants

As we found that there are differences in isoleucine and asparagine production by *S. aureus* strains from AD as compared to HC, we hypothesized that treatment of *S. aureus* and *S. hominis* strains with these amino acids would impact growth. A subset of the *Staphylococcus* strains was selected for growth curves analysis based on isoleucine and asparagine production levels and the amount of biofilm biomass formed (Table S14). The *S. aureus* strains from severe disease grew to a higher density than strains from healthy participants when grown in the presence 15 µM of isoleucine (Figure 4E). In contrast, while asparagine supplementation at 15 µM does impact strain growth in some isolates, with isolates HC08-R-ID-1 and AD34-L-AC-LS-1 exhibiting increased growth (Figure 4F), the effects are variable.

### Isoleucine and asparagine reduce S. hominis growth

Having shown earlier that there is an inverse relationship between *S. aureus* and *S. hominis* on AD skin, we next set out to test if the amino acids released by *S. aureus* during growth could impact *S. hominis* growth. In general, the addition of isoleucine (15mM) and asparagine (15mM) suppressed *S. hominis* growth, with the effect being more pronounced in strains from healthy participants (Figure 4G, Figure 4H). Interestingly, asparagine supplementation for the isolate AD42-L-AC-NLS-3 did not result in any observable influence on growth, reinforcing the variable impact of asparagine on staphylococcal growth (Figure 4H).

## Discussion

Although the key role of *Staphylococcus* species in AD has been well established, the role of health associated *Staphylococcus* species and the impact of strain variation in AD are yet unclear. In this study, we isolated staphylococcal strains from AD and HC participants in HK and analysed these strains using a combination of genomic and microbiological omics techniques. In doing so, we identify there are key differences in strains between disease and healthy participants. In this study, a well-controlled cohort of paediatric and adolescent participants from HK was recruited. Aside from topical corticosteroid application, concurrent skin disease, systemic medication, and personal cleaning practices were strictly controlled for to minimize confounding factors. Although it would have been ideal to also control for topical corticosteroid usage, this would not have been ethical or feasible as topical corticosteroids are the common first line treatment for AD. Further, while the usage of moderately potent or highly potent topical corticosteroids has been shown to reduce the density of *S. aureus* in AD, there is no direct evidence for low potency topical corticosteroids having an impact on the AD skin staphylococcal profile ^40^. As the skin staphylococcal profile generated from this study closely matches that of previous studies, this further reinforces the argument that there is minimal confounding from including patients on topical corticosteroids in the study ^5,9,25^.

In other studies, the *S. aureus* isolates collected from healthy participants have mainly come from the nares, as it is rare for *S. aureus* to be present on healthy skin ^14,17^. Here, the extensive efforts in culturing 835 staphylococcal isolates allowed us to ultimately collect the 16 *S. aureus* isolates from the matched skin site (non-nares) of 5 out of 24 healthy participants. This allowed us to directly compare the *S. aureus* strains present on healthy skin and diseased skin. Interestingly, each of the *S. aureus* strains found in healthy participants were also found in AD patients, suggesting that *S. aureus* colonization alone is insufficient to cause disease.

The investigation into the genomic background of *S. aureus* isolates found in AD has an extensive literature ^13,15–18^. While no single *S. aureus* genotype or clonal complex has been linked with AD, most AD patients are shown to be colonized stably by a single *S. aureus* genotype ^41^. In agreement with this finding, we made a similar observation in this study. However, as we had the AD severity scores of our participants, we were able to extend this finding beyond a binary disease vs health comparison, and show that with increasing severity, an individual carries more than a single *S. aureus* genotype. Notably, if multiple *S. aureus* genotypes were found in a patient, these genotypes belonged to the same *spa*-CC. This suggests that within each community, the multiple *S. aureus* strains evolved from a common ancestor. The selection pressures that may drive *S. aureus* microevolution on AD skin is an area for future exploration that will potentially provide further information for mechanisms of disease progression.

In addition to the genotypic analysis conducted, the phenotypic characterisation of strains here clearly delineates a cut-off between *S. aureus* strains found in mild and severe disease. Previous studies were unable to make such a demarcation as the comparisons were only between healthy and AD ^18,42^. Here, we provide an understanding of how *S. aureus* strains may colonize the skin prior to AD onset, with moderate and severe-associated strains growing faster than mild and health-associated strains. Once established on the skin, we found the *S. aureus* strains from severe AD formed more biofilm compared to strains from less severe AD. Interestingly, biofilms have previously been shown to be a virulence factor in AD by aiding persistence on the skin ^43^. Our observation that strains from severe AD had a lower carrying capacity than those from mild AD is important because this demonstrates that severity is not a simple function of bacterial abundance, but rather of bacterial functional activity. Thus, our novel findings of how *S. aureus* strain growth and biofilm formation are associated with disease severity suggests that investigating strains would help to identify key mechanisms involved in AD.

Building upon the findings for growth and biofilm formation, NMR was used to investigate the metabolome of *Staphylococcus* strains. This approach is unique, as previous metabolomic studies in AD have looked solely at the host production of metabolites and not the microbial production of metabolites ^44^. The discovery of isoleucine and asparagine being released at higher levels from *S. aureus* strains with increasing AD severity reinforces the concept that there is a clear difference between *S. aureus* strains found in mild and severe disease. Notably, experimental validation indicated that these two amino acids play a role in modulating phenotypic behaviors by staphylococcal species. While we observed isoleucine increasing growth in diseased *S. aureus* strains and suppressing growth in tested *S. hominis* strains during experimental validation, the fact that we did not see a similar consistent result for all strains for asparagine hints at strain specificity in this phenomenon.

Interestingly, isoleucine and asparagine have been studied before in AD, albeit with a limited understanding of the mechanism by which these amino acids affect disease. The topical application of L-isoleucine has previously been shown to alleviate facial AD ^45^. Likewise, a metabolomics study of skin biopsies found that were higher levels of asparagine on AD lesional skin as compared to AD non-lesional skin ^46^. While both of these studies present opposing evidence to our results, it brings about a salient point about the interpretation of metabolites. Metabolites are no longer considered to be exclusively used by other cells ^47^. Rather metabolites can also be used by cells they originated from in an autocrine fashion, indicating the role of metabolites as active players of cellular programs. For example, it may be that *S. aureus* or other microbes on the skin are using the isoleucine produced by the *S. aureus* strains, before it can be metabolized by the skin. Thus, a high level of isoleucine, the isoform of isoleucine, or the ratio of isoleucine to other metabolites in the system could result in the production of other factors that have an impact on skin. Notably, isoleucine can be metabolized into acetyl-CoA or propionyl-CoA, which are ultimately involved in the citric acid cycle and can be used by microbes for energy production ^48^. This may explain why we observed the increased growth of *S. aureus* when supplemented with isoleucine.

This study, despite the extensive work conducted on staphylococcal strains in AD, has some limitations. Firstly, the observed strain differences in growth, biofilm, and staphylococcal metabolome were made by growing the strains in a nutrient-rich media (TSB). To further advance the findings of the strain differences in growth, future work would be to look at the growth of the strains using different media, such as minimal media and artificial sebum. Secondly, the experimental validation of isoleucine and asparagine suggests that these two amino acids may be a means by which *S. aureus* can establish its niche in the skin microbiome by impacting on the balance between *S. aureus* and commensals such as *S. hominis*. The next step would be to investigate the mechanism by which isoleucine and asparagine inhibit *S. hominis* growth. Thirdly, this work notably focused on the microbe (*S. aureus* and *S. hominis* respectively) and potential microbe-microbe interaction (*S. aureus – S. hominis*), being the first study to explore these interactions in AD-associated strains. To build further upon this work, the next step is to introduce the microbes into models that represent the host: mock skin community and epithelial cells/epidermal layer models, as well as murine models of AD.

In this study, we demonstrate that specific staphylococcal strains with distinct phenotypic behaviours are associated with AD disease severity. We further show that this association may be mediated by isoleucine and asparagine production by *S. aureus*. In doing so, we provide a potential mechanistic link of how the skin microbiome could impact disease severity in individuals that develop AD. As such, we identify novel areas for therapeutic intervention in the management of this inflammatory skin condition.

## Data availability

All data and code needed to evaluate and reproduce the results in the paper are present in the paper and/or the Supplementary Figures and Tables.

## Acknowledgement

We thank all the families that took part in this study and the support from the nurses in the Department of Pediatrics and Adolescent Medicine and Division of Dermatology at the Queen Mary Hospital in Hong Kong.

## Author contributions

D.L.M. and H.M.T. conceived and designed the study. G.T.C., C.S.M.W., and J.S.R.D recruited participants. H.K.F.W., V.C., and J.Z. extracted DNA from the bacterial isolates and conducted *spa* typing of the *S. aureus* isolates. H.K.F.W, X.Y.W., and C.C.Y.L performed the growth curves assay. H.K.F.W. and X.Y.W performed the biofilm crystal violet assay. H.K.F.W. and S.K. prepared the exhausted media for NMR metabolomics. A.L.G. performed the NMR metabolomics and peak picking for the metabolite identification. H.K.F.W. carried out the data and statistical analyses. A.N.P., M.Z.J., J.M.Z., E.S.H, Y.C., H.A., and E.D. reviewed and edited the manuscript. H.K.F.W., D.L.M., and H.M.T. wrote the manuscript with input from all co-authors. D.L.M. and H.M.T acts as the guarantor for this study and publication.

