## Supplemental Files for "Isoleucine and asparagine influence the growth of staphylococcal strains in atopic dermatitis"

**Supplementary Figures**

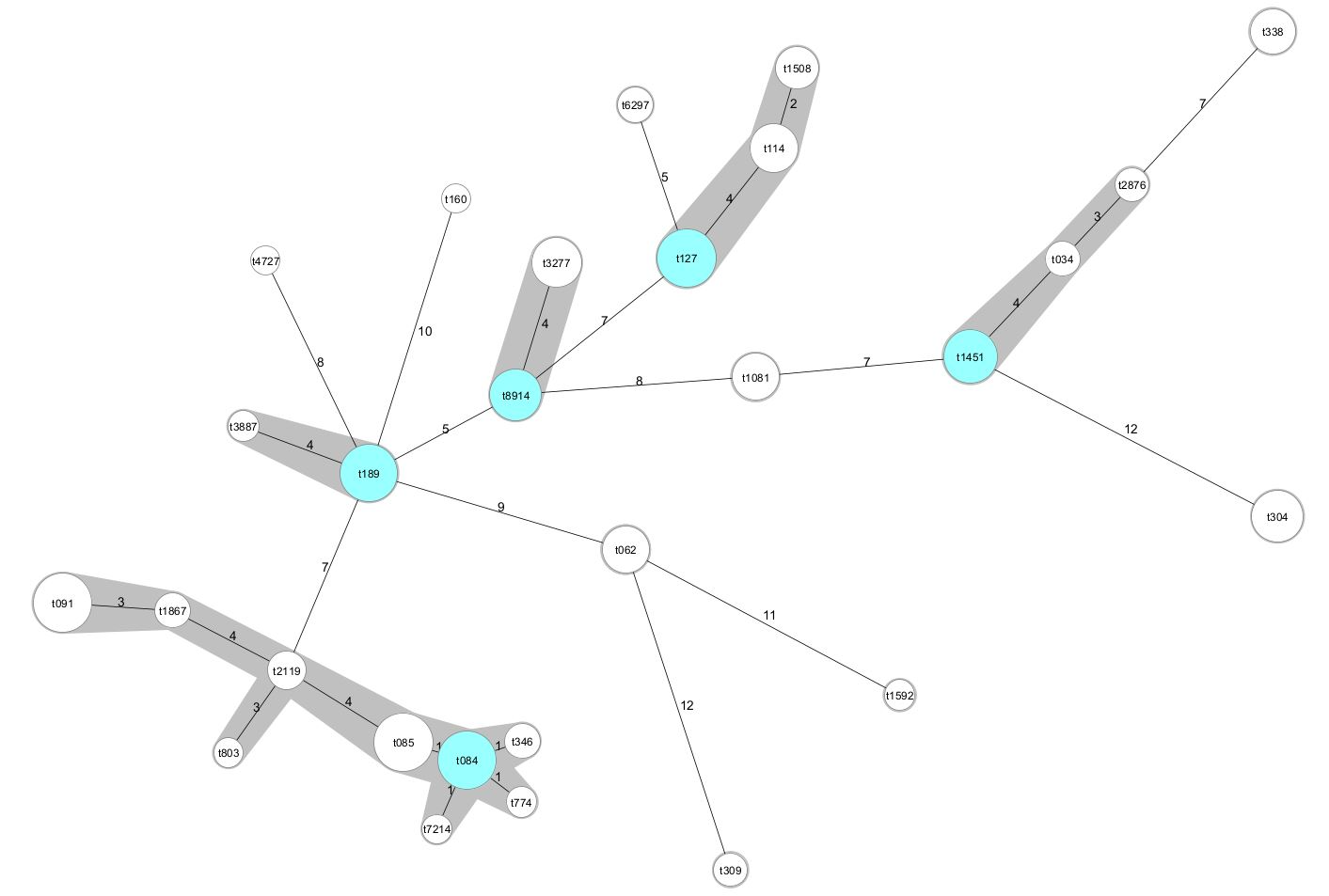
**Figure S1. Cluster analysis of *spa* types.**

The BURP clustering algorithm was used to assign *spa* types into a *spa* clonal complex (*spa*-CC) (colored in dark grey). Each circle represents a *spa* type, with the size of the circle being proportional to the frequency for a given *spa* type. Circles in blue are the founder *spa* types for a *spa*-CC. Circles in white are member *spa* types for a *spa*-CC. The number on the lines connecting two *spa* types represents the number of repeats in the polymorphic X region that differ between the two *spa* types.

**Supplementary Tables**

**Table S1. Subject inclusion and exclusion criteria.**

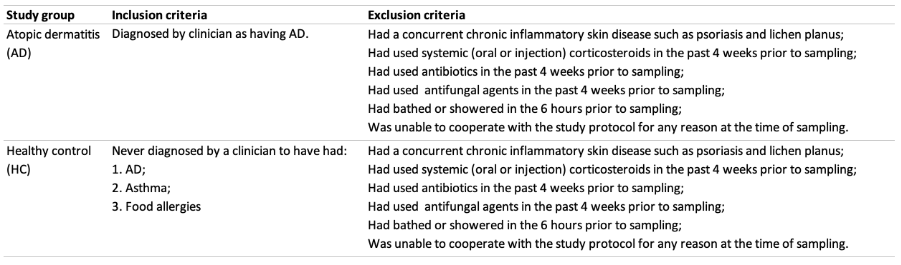

**Table S2. Count of *Staphylococcus* species isolates cultured by severity.**

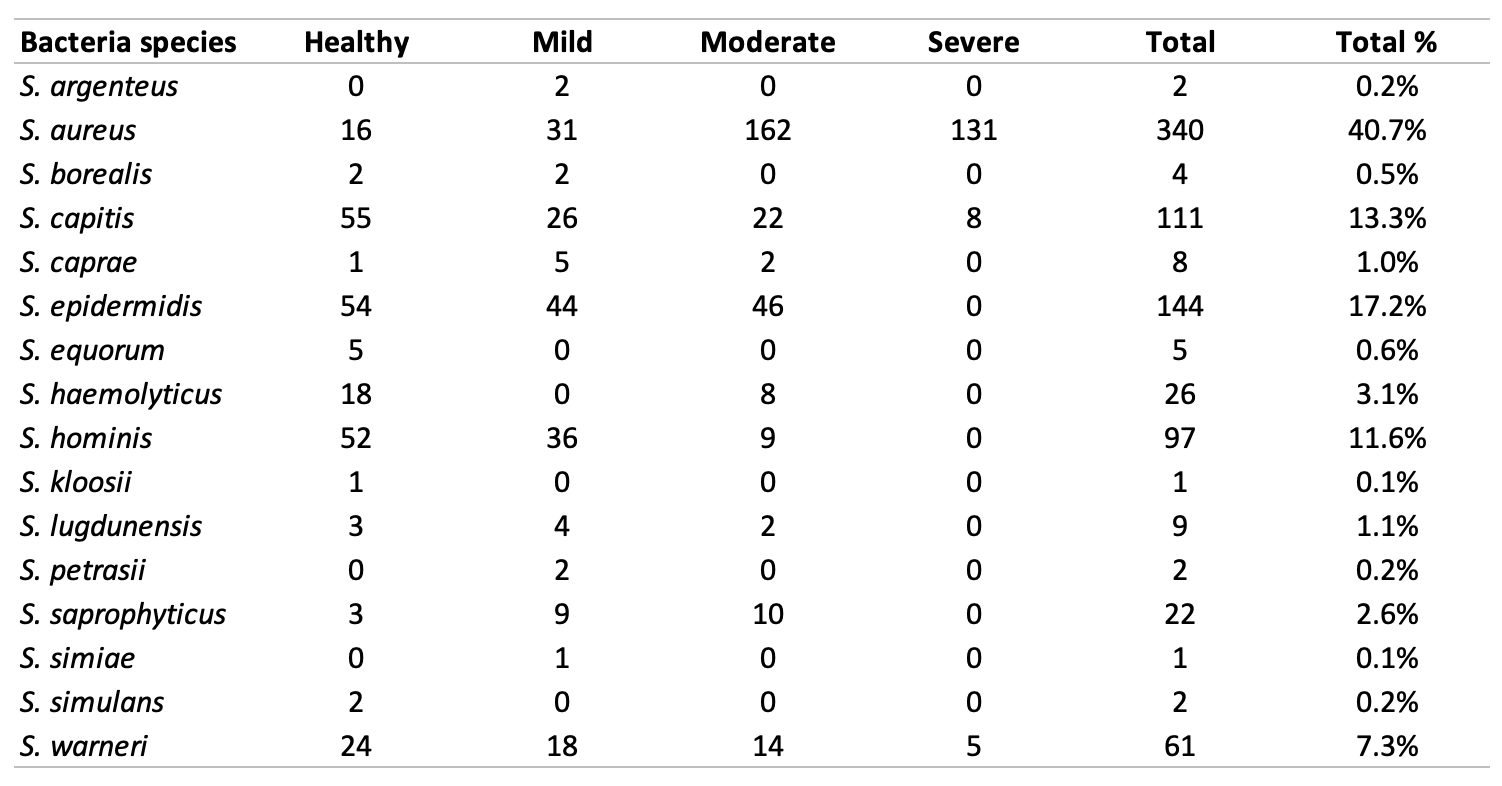

**Table S3. Presence and absence of *Staphylococcus* species isolated by severity.**

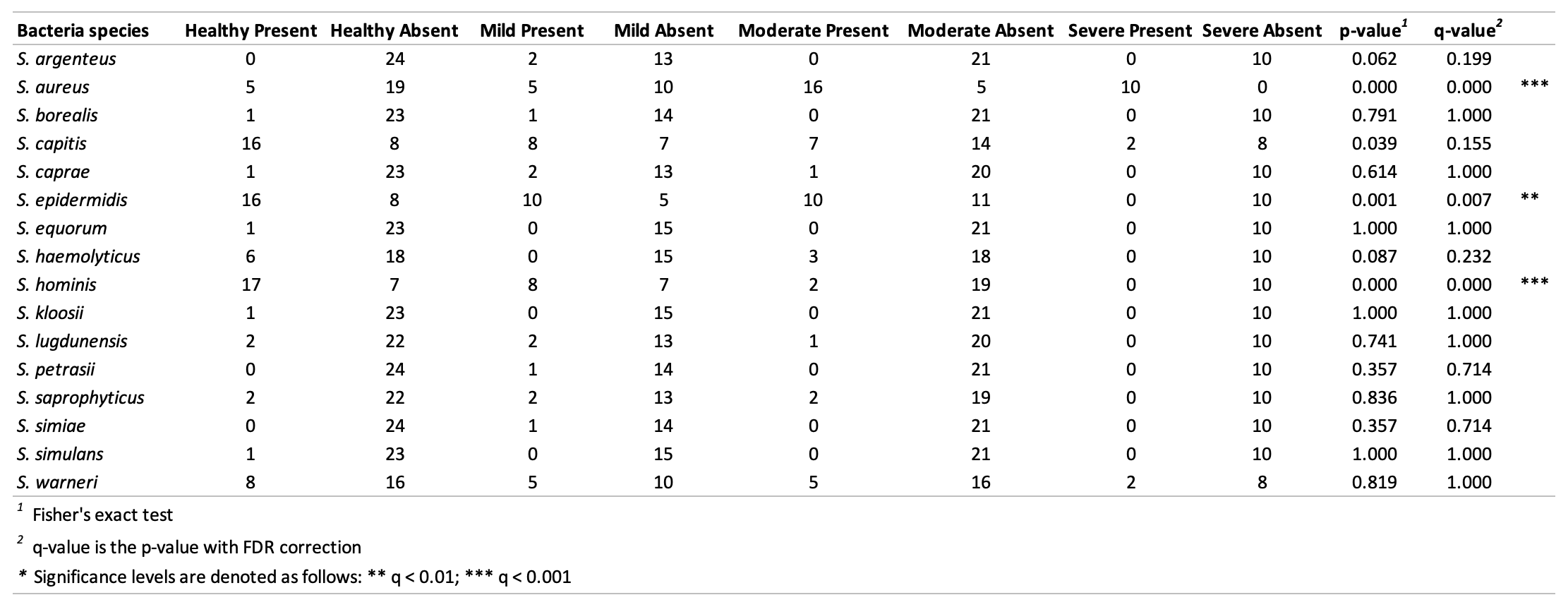

**Table S4. Percentage of subjects by severity having a *Staphylococcus* species on their skin.**

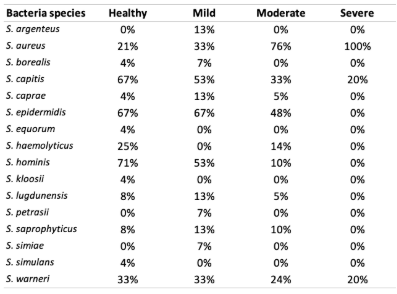

**Table S5. Distribution of *S. aureus* genotypes isolated from AD and HC subjects.**

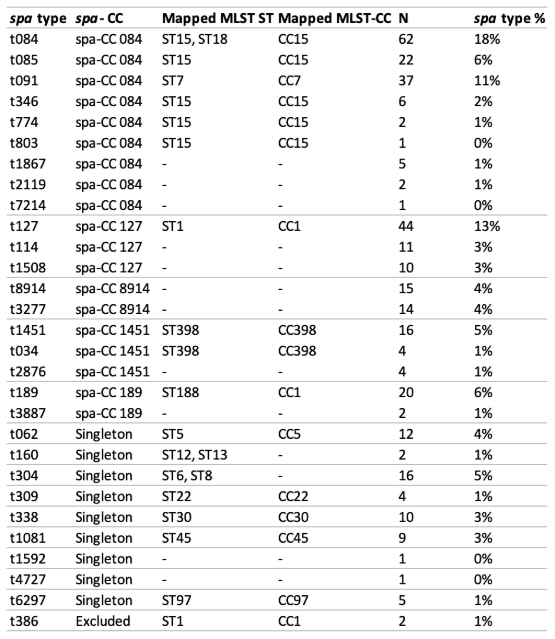

**Table S6. Relationship between *spa* types and disease status.**

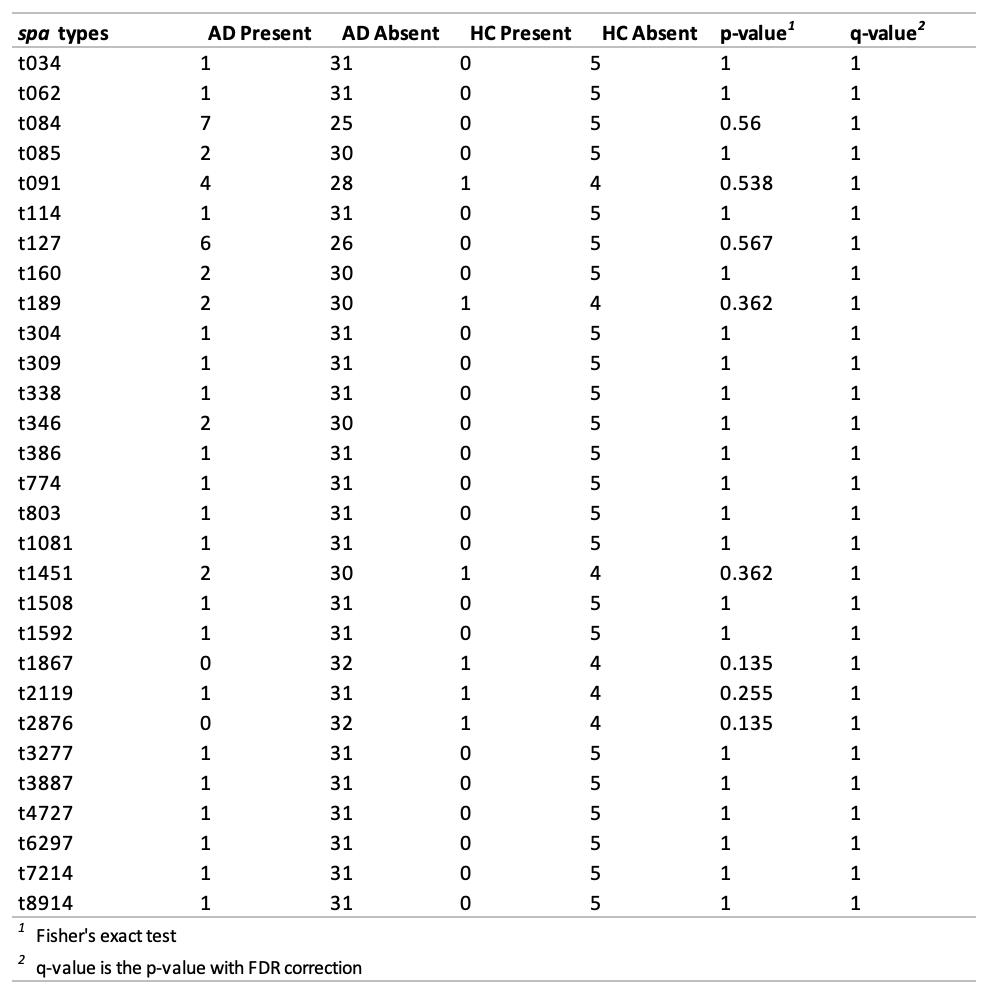

**Table S7. Relationship between *spa*-CCs and disease status.**

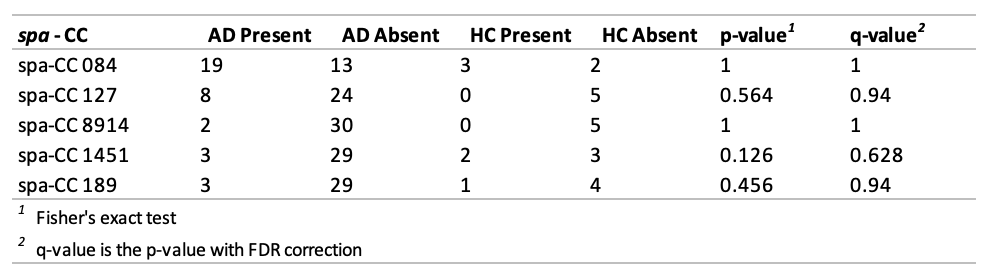

**Table S8. Relationship between MLST STs and disease status.**

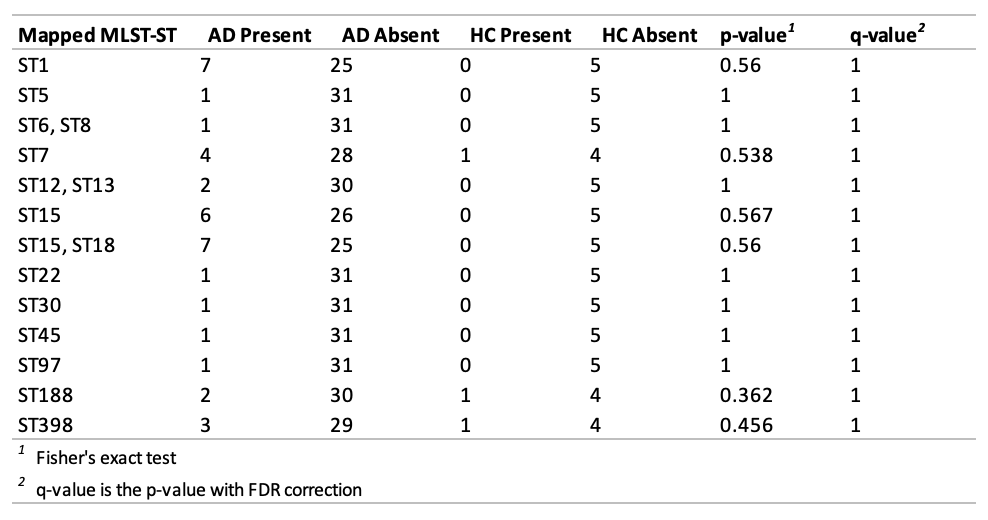

**Table S9. Relationship between MLST-CCs and disease status.**

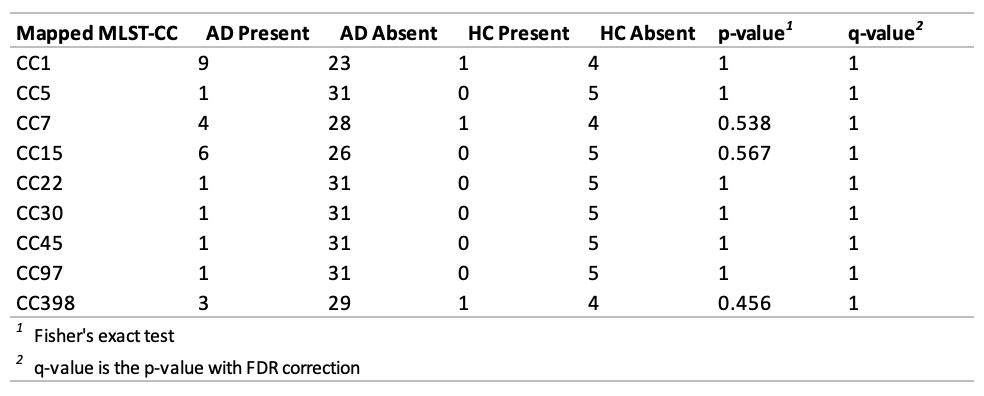

**Table S10. *S. aureus* isolates picked for onward analysis.**

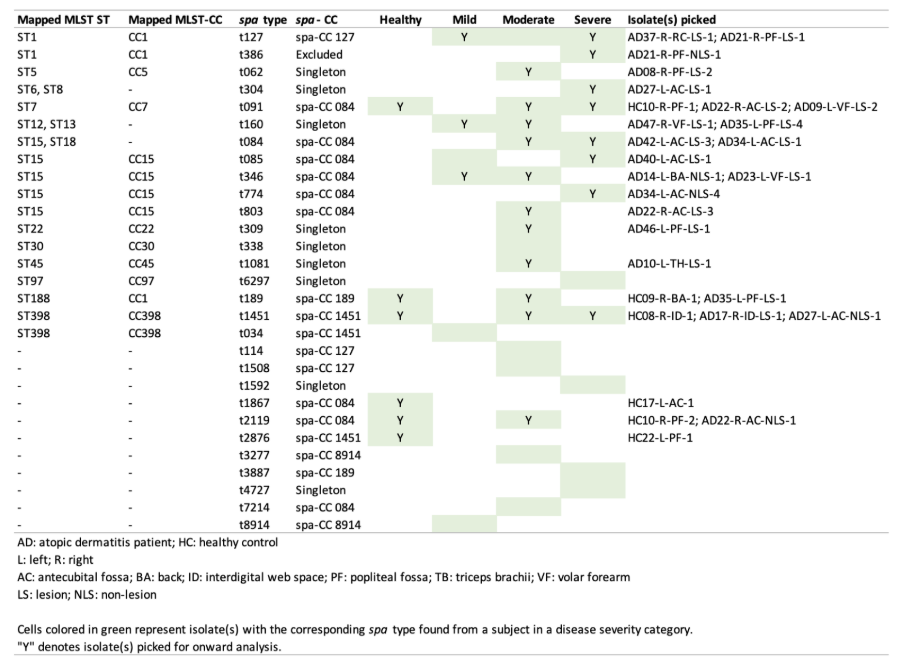

**Table S11. *S. hominis* isolates picked for onward analysis.**

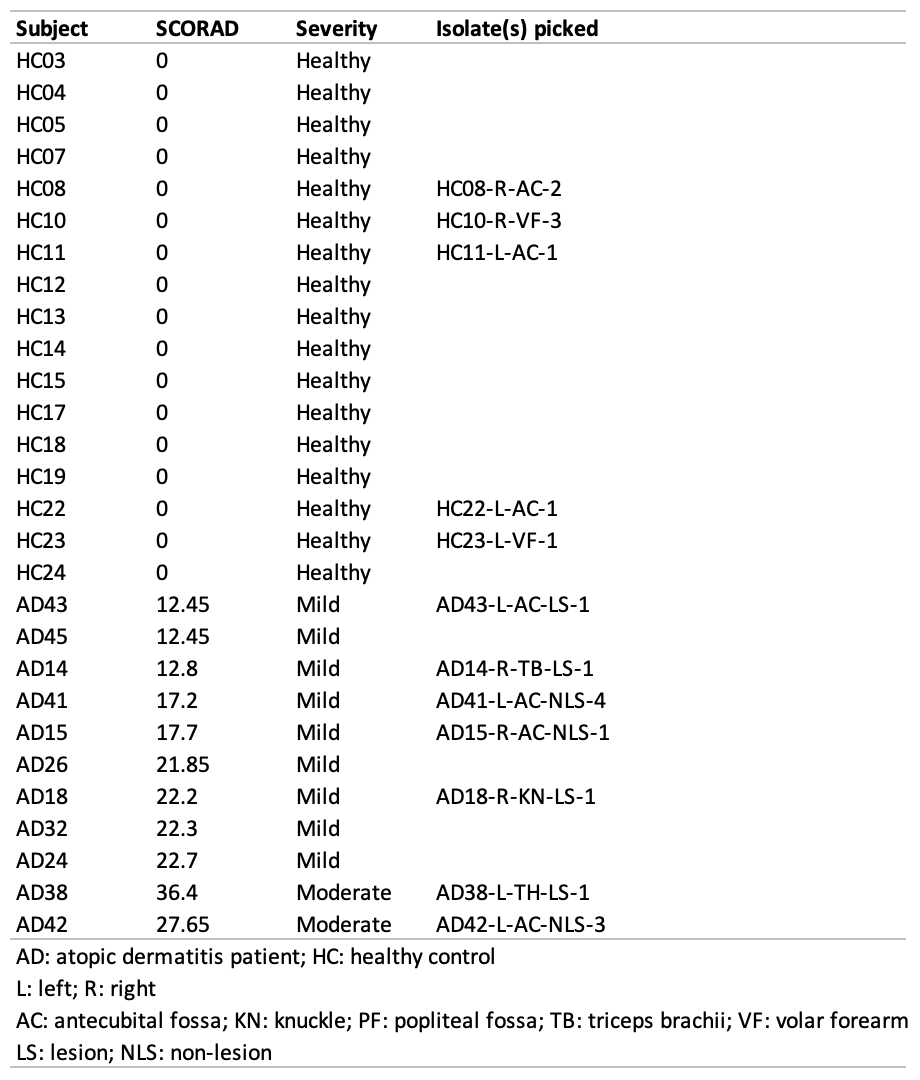

**Table S12. Metabolites identified from NMR metabolomics in *Staphylococcus* strains.**

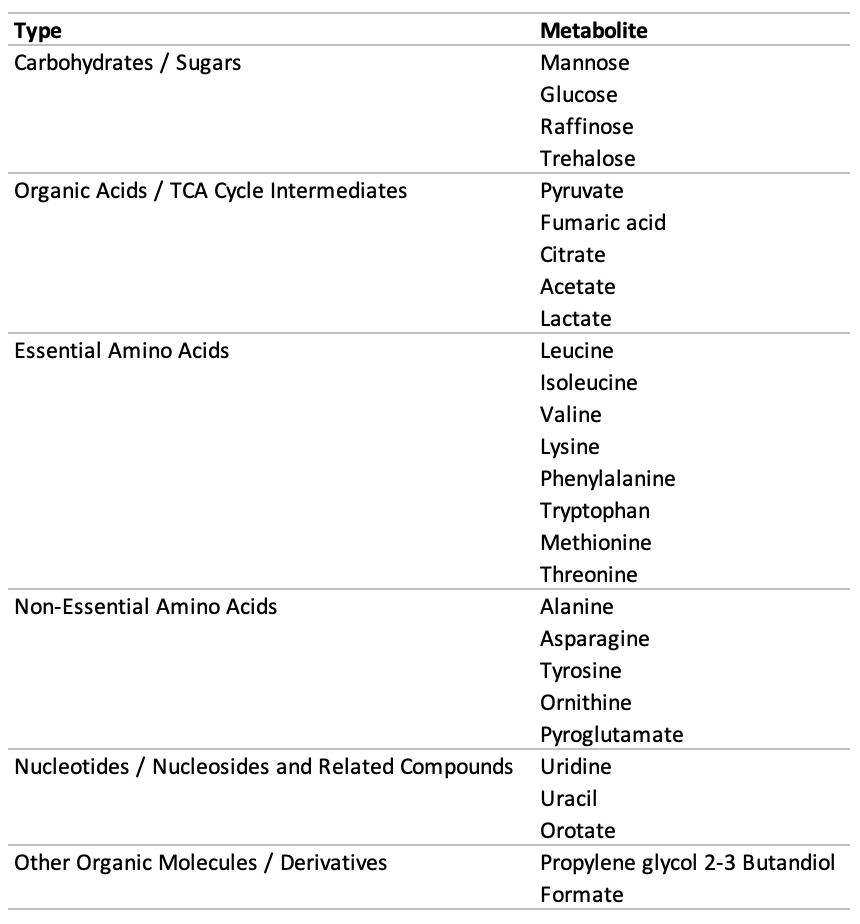

**Table S13**. **Distribution of metabolites produced by *S. aureus* strains from AD and HC.**

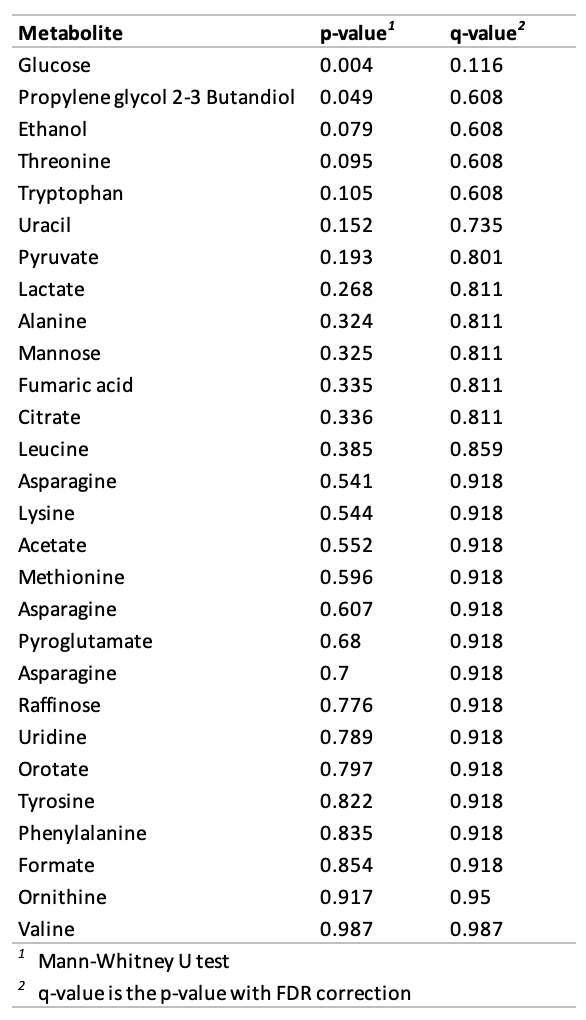

**Table S14. *Staphylococcus* strains selected for growth analysis with isoleucine and asparagine treatment.**

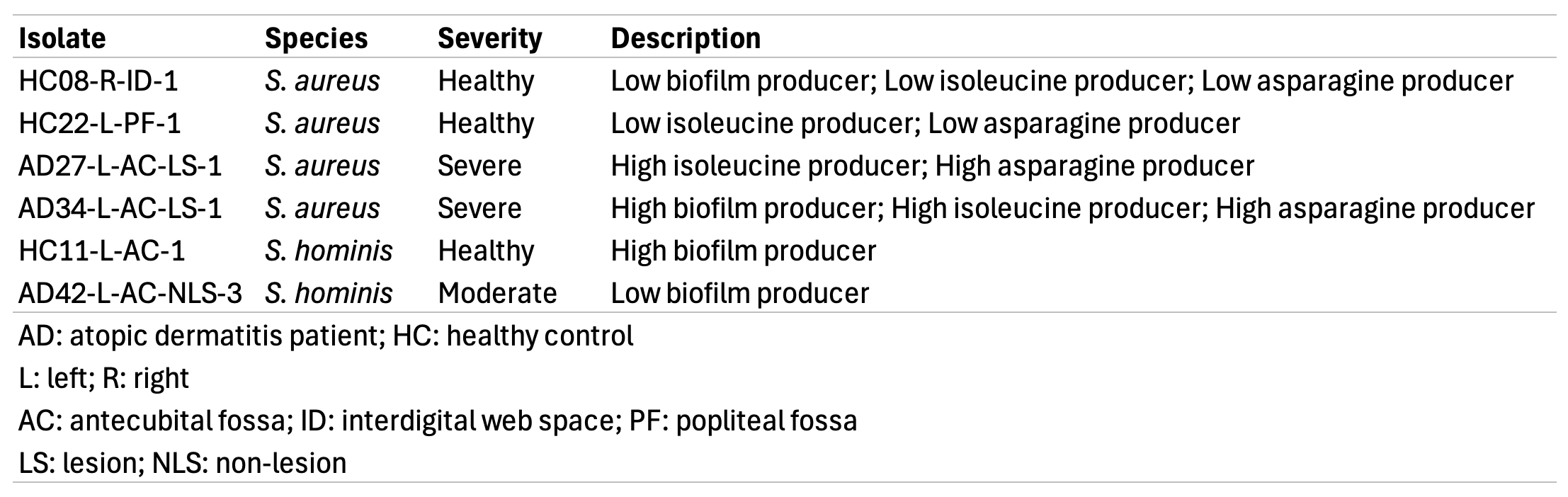
